# Rotamer libraries for the high-resolution design of β-amino acid foldamers

**DOI:** 10.1101/086389

**Authors:** Andrew M. Watkins, Timothy W. Craven, P. Douglas Renfrew, Paramjit S. Arora, Richard Bonneau

## Abstract

β-amino acids offer attractive opportunities to develop biologically active peptidomimetics, either employed alone or in conjunction with natural α-amino acids. Owing to their potential for unique conformational preferences that deviate considerably from α-peptide geometries, β-amino acids greatly expand the possible chemistries and physical properties available to polyamide foldamers. Complete *in silico* support for designing new molecules incorporating nonnatural amino acids typically requires representing their side chain conformations as sets of discrete rotamers for model refinement and sequence optimization. Such rotamer libraries are key components of several state of the art design frameworks. Here we report the development, incorporation in to the Rosetta macromolecular modeling suite, and validation of rotamer libraries for β^3^-amino acids.

## 1. Introduction

The explicit treatment of the conformation of side-chain atoms distinguishes low-resolution and high-resolution models of protein structure.[1] A common low-resolution approach is to instead approximate the side chain with a single pseudoatom located at the center of mass;[2] such methods require less physically realistic energy functions[3, 4] but permit faster sampling over a smoother energy landscape.[5, 6] Explicit (all atom) side chain conformations may instead be efficiently sampled using a rotameric approximation, where discrete sets of side chain χ angles are chosen from well-validated conformations that are meant, ideally, to correspond to the conformational energy minima for the side chain in question.[7]

These sets of side chain conformations are called ‘rotamer libraries.’ Sampling side chain conformations from rotamer libraries is especially efficient when there is substantial interdependence between side chain degrees of freedom, and when only a small fraction of the plausible local minima are near the global minimum. Rotamer sampling requires rotamer libraries from which to sample conformations and recent works have demonstrated that these rotamer libraries should be dependant (to further improve computational efficiency) on backbone conformation.[8] Prior development of noncanonical rotamer libraries for use in Rosetta has employed side chain coordinate scans and energy optimization, either through quantum mechanics software or a molecular mechanics force field, followed by *k*-means clustering to generate rotamer libraries for NCAAs and peptoids.[9]

For canonical α-branched amino acids, the Dunbrack and Richardson labs have curated the most widely used rotamer libraries. In both cases, these libraries are derived from a statistical treatment of high quality experimental structures.[10–15] The Daggett lab has recently taken an alternate approach and employed molecular dynamics simulations[16, 17]. Work on these MD-based rotamer libraries points to several limitations to statistics-derived rotamers and also demonstrates that simulations can model side chain configurational clusters and minima when statistical methods are not applicable, as in the case of NCAAs.[18, 19]. Both this work and prior work by Kuhlman and Renfrew suggest that there exists a valuable role for rotamer libraries derived from simulations as well as experimental structures. Furthermore, since structural data on non-natural residues is very limited, general rotamer libraries for these moieties must be simulation-derived.

Peptides entirely composed of or containing β-amino acids (Figure 1) possess useful pharmacological properties, particularly slow proteolytic degradation and a curbed immune response.[20] Such molecules have been used to mimic apolipoprotein[21] and as antimicrobial compounds;[22] they may target mdm2,[23] the GLP-1 receptor,[24] or HIV fusion.[25] Detailed efforts to target BCL-2 family proteins[26] and GLP-1[27] have been described.

**Figure 1:**
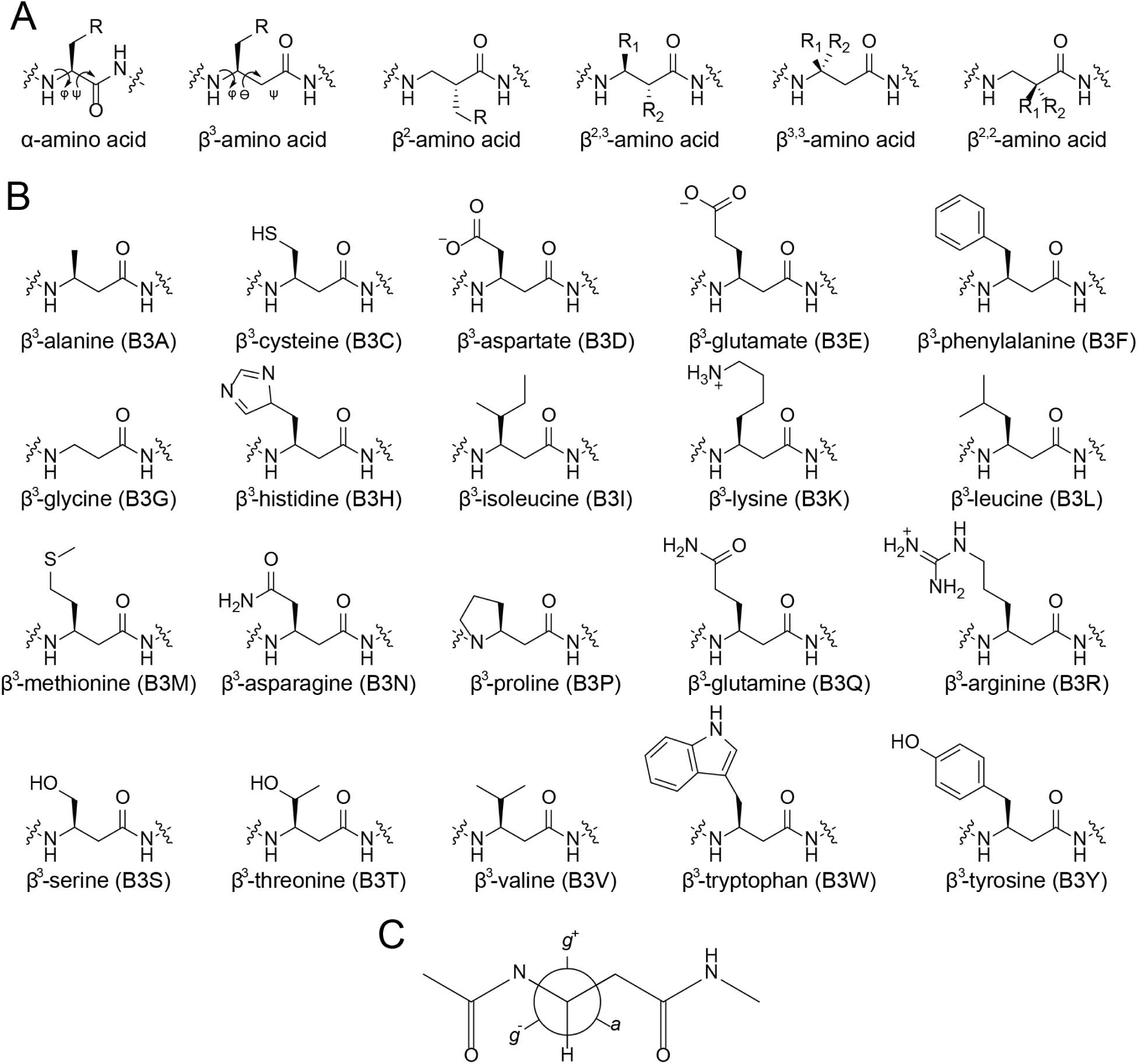
Schemes of chemical species referred to in the text. A. The α-amino acid and β-amino acid are marked with two mainchain torsions that were considered equivalent in prior representations of β-amino acids in Rosetta. While this work only considers rotamer libraries for β^3^-amino acids, the expansions to the Rosetta code enables representation of any β-amino acid or, indeed, any polyamide, using the same framework. B. The twenty β^3^-amino acid side chains, eighteen of which are described with rotamer libraries developed in this study. β^3^-glycine and β^3^-alanine do not possess side chain degrees of freedom. C. The **g**^+^, **g^-^**, and **a** conformations described here, for χ^1^: N-Cβ-Cγ-X dihedrals of 60°,−60°, and 180°.

These biological applications are enabled by the conformational preferences of β-amino acids. Their two backbone carbons provide four stereoisomers and six disubstituted isomers, leading to considerable conformational diversity. Helical structures achievable by β^3^-amino acids have been extensively studied. These include either the 3_14_ or 14-helix, characterized by a helical axis opposite the α-helix and a 14-atom hydrogen bonding macrocycle, and the 2.5_12_ or 12-helix, whose handedness is like the α-helix.[28,29] Helical bundles similar to α-helical α-amino acid coiled coils are accessible to 3_14_ helices, as well,[30] and one has been designed to have catalytic activity.[31]

β-amino acids also form local structures that are ideal for mimicking parallel[32–34] or antiparallel sheets.[33, 35–37] Such extended conformations may be adopted by cyclic β-peptide oligomers, but steric effects from mutliple substitutions can provide a similar conformational bias. For example, similar extended conformations are favored by *trans*-2,3-disubstituted residues by encouraging an *anti θ*. In contrast, 2,3-disubstituted cyclic sidechains stabilize *gauche θ* conformers that favor turns and helices,[37, 38] while achiral β^2,2^ or β^3,3^-disubstituted residues stabilize reverse turns.[39]

Though β-amino acids can form diverse conformations on their own, the accessible chemical space may be expanded further by using them in chimeric peptides, i.e. in combination with α-amino acids. Replacing an α-amino acid with a β-amino acid on both strands of a β-hairpin extends the length of the hairpin by one methylene unit but maintains the hydrogen bond network and so only tertiary contacts must adjust to accommodate the perturbation.[40, 41]

Chimeric α/β peptides can adopt helical conformations.[42–46] Optimized compositions ultimately involved organizing all the β-residues on a single helical face by employing β^3^-amino acids every third or fourth position.[47]

Due to the high therapeutic value of mimicking an α-helix[48–51] and the conceptual simplicity of homo-oligomers, peptides composed of or containing β-amino acids have been applied primarily as α-helix mimetics.[52] Consequently, β-amino acid rotamer libraries have been developed only for all-β conformations that form helical folds. There are high resolution rotamer libraries for β^3^ and β^2^ amino acids available for individual backbone conformations: those of 3_14_ and 3_12_ helices.[53] Such rotamer libraries, while valuable, are not suited to a general peptidomimetic context in two ways. First, they do not cover backbone conformations that are entirely favorable for β-amino acids but do not form homopolymer secondary structures, and so they are limited in application. Second, rotamer libraries fit to the rotamers assumed by an entire secondary structure element do not necessarily apply to other polymeric contexts. For example, the dihedrals assumed by a β-amino acid in a 3_14_ helix are only 20° from those assumed by a β-amino acid in an α3β helix, but the α3β helix has an opposite dipole moment and each β-residue is flanked by two α-residues. The resulting bias on rotamer libraries can be substantial; long range effects from adjacent helical turns are known to bias PDB-based rotamer libraries for at least valine, leucine, and isoleucine.[19] As a concrete example, the (**a, g^+^**) rotamer common in α-helical α-leucine residues creates no clashes in the all-β3_14_-helix depicted in Figure 2, but is incompatible with a common conformation of the chimeric α3β helix. Thus, in order to apply to a general peptidomimetic context, rotamer libraries must be constructed independent of non-local influences.

**Figure 2:**
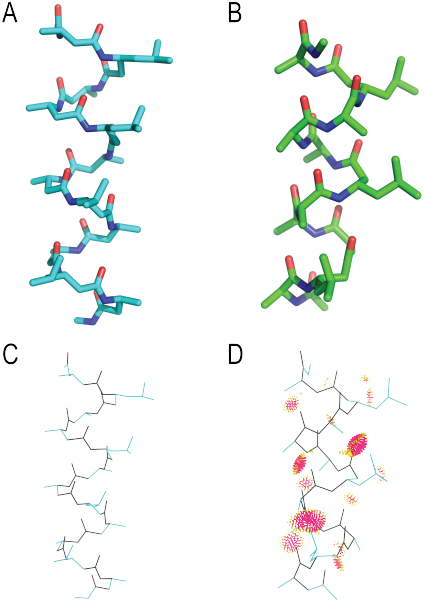
The (**a, g**) rotamer imposed on every fourth residue (a leucine) in the 3_14_ helix (A) and the α3βhelix (B). Molprobity clash analysis reveals that the 3_14_ helix is free of clashes (C), while the α3βhelix (D) has a clashscore of 114, far worse, in 12 residues, than low-quality structures of entire ribosomes. The distinct clash, shown in red dots, is with the *i* + 3 alanine methyl group.

These prior rotamer libraries have not been available for use in macromolecular modeling programs such as Rosetta, which has long assumed that rotamer libraries were dependent only on two mainchain dihedrals.[54] Prior work that attempted to model β^3^-peptides in Rosetta instead employed a limiting approximation: that the first two mainchain torsions of a β^3^-amino acid could be treated analogously to the φ and ψ torsions of an α-amino acid (Figure 1).[55] Backbone dihedral angles typical for a 3_14_ helix such as (−140°, 60°, −120°) are described by a high-energy of Ramachandran space like (−140°, 60°) and as a result are drastically undersampled.

Here, we have developed and incorporated into Rosetta backbone-dependent rotamer libraries for β^3^-amino acids. Furthermore, we generalized Rosetta’s treatment of rotamers and rotamer libraries such that it can create and use rotamer libraries with any number of backbone dihedral angles. We validated this approach both computationally and experimentally. Our results are in good agreement with quantum mechanical calculations on several side chains, which demonstrate that we recover the rotamer wells and their approximate populations. Our results also recover the rotamers found in high quality crystal structures in the Protein Data Bank. Furthermore, use of these rotamer libraries in conjunction with Rosetta recovers trends in experimental binding affinity in experiments with β-peptide helices. We anticipate that this rotamer library system will constitute an important step towards the direct, accurate rotamer-based design of structures containing arbitrary oligomers in Rosetta.

## 2. Results and Discussion

### 2.1. β-amino acids exhibit diverse backbone conformations

To quantify the need for a general, fully backbone dependent rotamer library for β^3^-amino acids, we explored the backbone conformations that they can adopt. Rather than setting an energetic threshold and exploring candidate energy functions, we simply sought out conformations that lacked intramolecular clashes. Table 1 compares the volume of backbone dihedral space that each β side chain may adopt without clashes versus its α amino acid equivalent. The volume within 40° of the 3_14_ helix or α3β helix dihedral angles is only 1.1% of the total backbone dihedral space of a β^3^-amino acid, illustrating that β^3^-amino acids may adopt far more than the two homopolymer conformations that have received most attention.

**Table 1:**
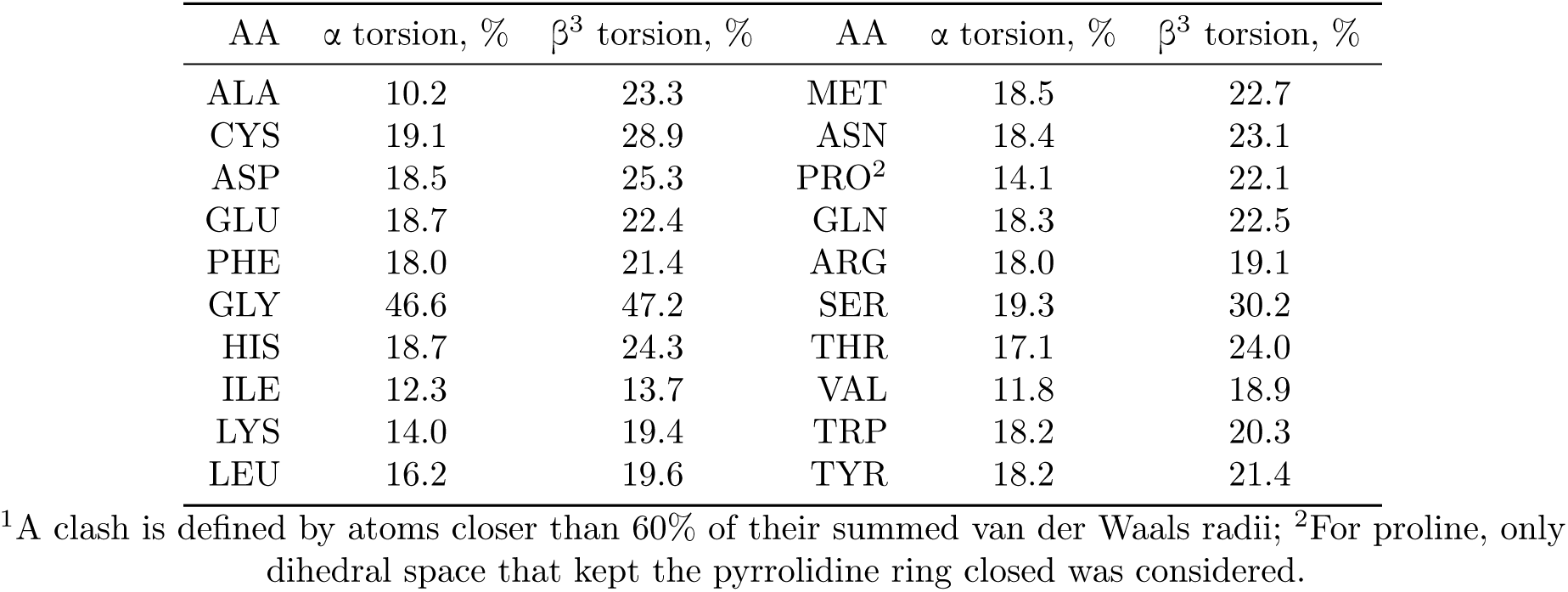
Proportion of backbone dihedral space (“Ramachandran space” for α-amino acids) that does not result in any clashes^1^

| AA | $\alpha$ torsion, % | $\beta^3$ torsion, % | AA | $\alpha$ torsion, % | $\beta^3$ torsion, % |
| --- | --- | --- | --- | --- | --- |
| ALA | 10.2 | 23.3 | MET | 18.5 | 22.7 |
| CYS | 19.1 | 28.9 | ASN | 18.4 | 23.1 |
| ASP | 18.5 | 25.3 | PRO<sup>2</sup> | 14.1 | 22.1 |
| GLU | 18.7 | 22.4 | GLN | 18.3 | 22.5 |
| PHE | 18.0 | 21.4 | ARG | 18.0 | 19.1 |
| GLY | 46.6 | 47.2 | SER | 19.3 | 30.2 |
| HIS | 18.7 | 24.3 | THR | 17.1 | 24.0 |
| ILE | 12.3 | 13.7 | VAL | 11.8 | 18.9 |
| LYS | 14.0 | 19.4 | TRP | 18.2 | 20.3 |
| LEU | 16.2 | 19.6 | TYR | 18.2 | 21.4 |

<sup>1</sup>A clash is defined by atoms closer than 60% of their summed van der Waals radii; <sup>2</sup>For proline, only dihedral space that kept the pyrrolidine ring closed was considered.

The generous standard of clash-free conformations is useful to assess the intrinsic flexibilty of β-amino acids, in contrast to homopolymer-preferred conformations, which might require regular, repeating stabilizing interactions. These conformations are important to multiple β-amino acid use cases, such as the design of substituted macrocycles, where the cyclic constraint enforces the local conformation and there is no requirement for regularity. In addition to the well-studied helical conformations, in the process of this investigation we found a range of extended conformations available to β-amino acids, which have not been previously investigated in any systematic way. Such extended conformations possess φ between −130° and −180°, with θ and ψ positive and summing to 30° to 50° less than |φ|.

### 2.2. The MM energy function reproduces the energy wells of quantum mechanics studies

The molecular mechanics scoring function employed in Rosetta, which we used to generate these rotamer libraries, reproduces the minima identified by quantum mechanical calculations. We evaluated four common β^3^-amino acid side chains: β^3^W, β^3^L, β^3^I, and β^3^S, restrained to two backbone conformations: a helical conformation reminiscent of both the 3_14_ helix, (−140°, 60°,−120°), and the α3β helix and an extended conformation that would form a pleated sheet, (−140°, 75°, 65°). We conducted constrained geometry optimization followed by single-point energy calculation at the β^3^LYP level of theory on side chain conformations sampled every 10°. The resulting energy surface as a function of χ1 and χ2 compares favorably to the molecular mechanics derived energy surfaces for each residue type, showing that we capture the same rotamers as the considerably more computationally expensive method (Figure 3). In order to quantify the accuracy with which the two heat maps agree, we computed the Bhattacharyya distance for each residue and backbone conformation (Table 2). The results verify the significance of the similarity of the energy landscapes, improving confidence in our results.

**Figure 3:**
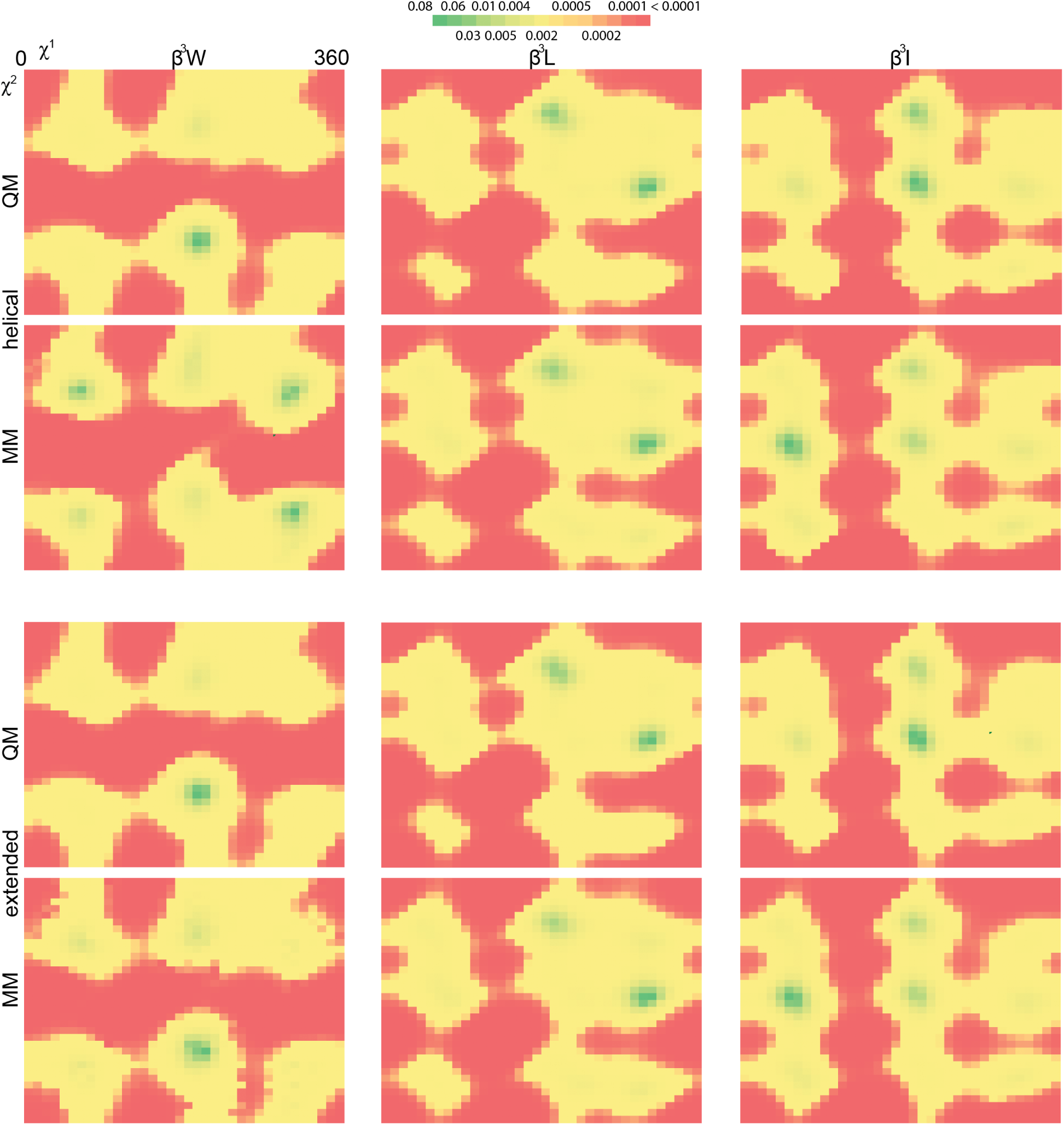
The probability distributions as determined by quantum mechanics (top) and molecular mechanics (bottom) for χ_1_(horizontal axis) and χ_2_ (vertical axis) for three residues (β^3^I, β^3^L, β^3^W; columns) in two conformations (helical and extended).

**Table 2:** The Bhattacharyya distance for each of eight pairs of probability distributions, representing the side chain conformational preferences of four β^3^-amino acids in two conformations, evaluated either at the quantum mechanical (QM) or molecular mechanics (MM) level of theory.

| AA | QM-MM Distance |  |
| --- | --- | --- |
|  | Helical | Extended |
| $\beta^3$I | 0.155 | 0.163 |
| $\beta^3$L | 0.048 | 0.069 |
| $\beta^3$S | 0.237 | 0.233 |
| $\beta^3$W | 0.261 | 0.074 |

### 2.3. The new β^3^-amino acid rotamer libraries reproduce observed PDB rotamers

A considerable number of structures containing β^3^-amino acids already exist in the Protein Data Bank, though all but one is in a mixed-α/β or 3_14_ helix. We built the same rotamer set as employed in Rosetta’s packing algorithm, by performing tricubic interpolation between adjacent backbone dihedral bins (i.e. the eight vertices of the cube of grid points for which explicit rotamers were solved). The resulting rotamer sets possess a rotamer within 15.5° of the experimental rotamer or better, with many closer than 10° (Figure 4).

**Figure 4:**
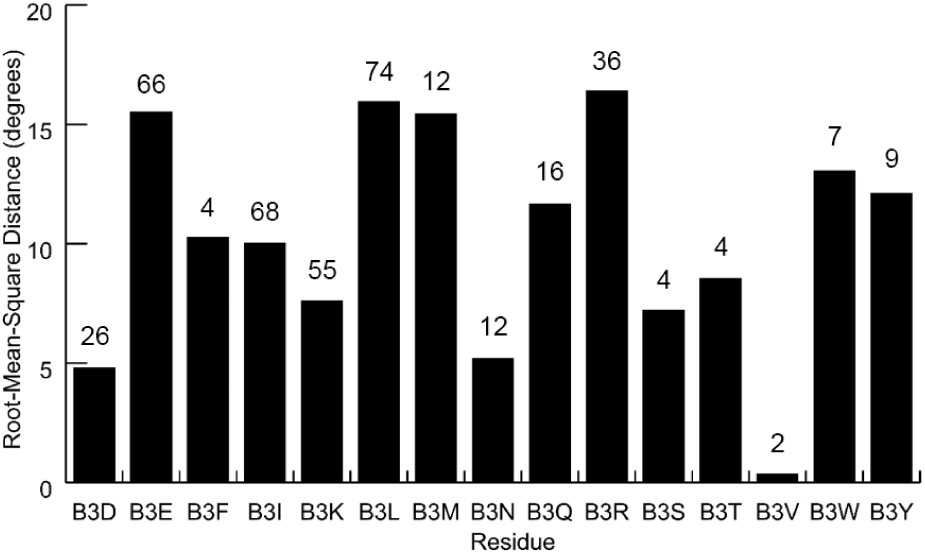
RMS error in degrees for nearest rotamer across all resolved β^3^-amino acid samples in the Protein Data Bank. No data exists on β^3^-homocysteine or β^3^-homohistidine, and rotamer libraries do not apply to β^3^-homoalanine or β^3^-homoglycine. Errors are comparable to those found in prior work on fitting peptoid and non-canonical amino acid rotamer libraries[1, 9] and to the effective uncertainty of Dunbrack rotamer libraries (see Supporting Information).

This performance in native rotamer recovery is comparable to the performance of the Dunbrack libraries on the equivalent α-amino acid side chain (see Supporting Information); especially considering the narrow scope of the test data only one region of backbone torsion space) against the general model employed here, we find this result very encouraging.

In the process of assessing the errors in PDB rotamer recovery, we found that several of the native rotamers we had been treating as ‘gold standards’ inhabited structurally improbable conformations. Since they were not forming tightly packed interactions in a hydrophobic core, nor with a binding partner, nor with a crystal contact, we suspected that they might constitute structural errors. We examined several representative structures that possessed high-energy native rotamers and built maps from the publicly available structure factors; in doing so, we found that many β-amino acid residues with very little electron density supporting any particular side chain conformation; for example, we highlight four examples from 3G7A and 4BPI in Figure 5, but others are found in 4KGR, 3O42, and more. We strongly suspect that many of the deviations observed here are due to rotamers that were built poorly in the absence of electron density constraints; these experimental samples are thus a poor reflection of physical ground truth.

**Figure 5:**
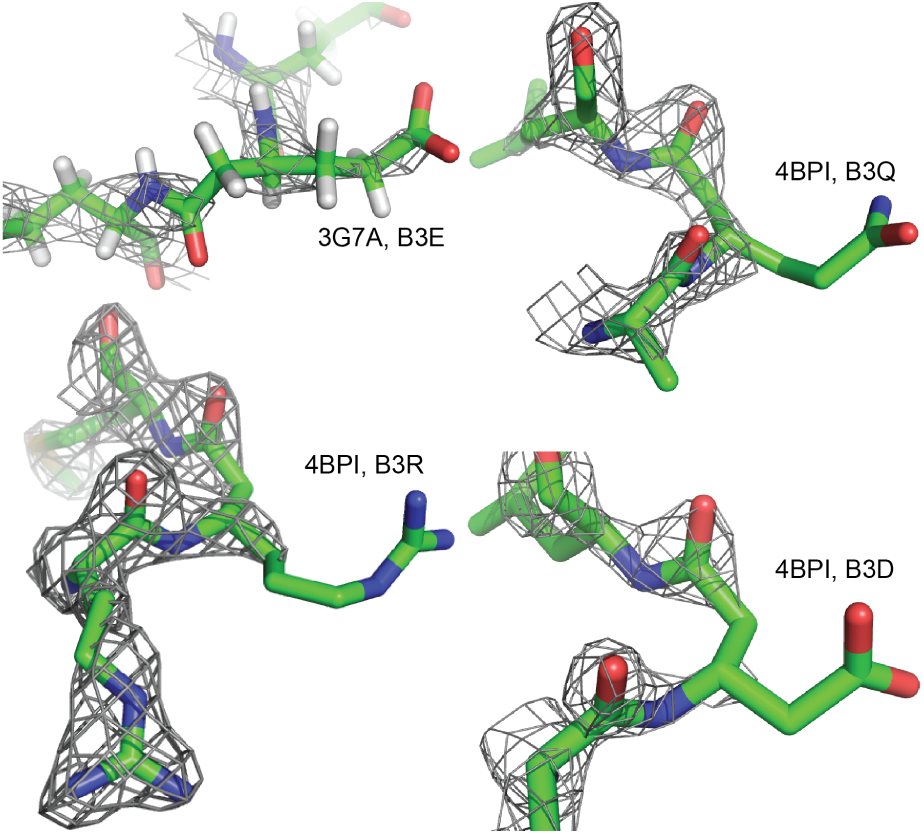
Often, our ‘missed’ side chains for a given β-amino acid appeared to have very high-energy native rotamers in the crystal structure. Due to the challenges of fitting atomic coordinates to electron density, and due to the additional challenge of phasing diffraction data for novel heteropolymers, structural data might be particularly uninformative for these molecules. Indeed, maps frequently provided minimal density near β-amino acid residues, as in the four examples above (PDB ID: 3G7A, 4BPI).

Since the total number of β-amino acid rotamers in the PDB is so small, we cannot perform this analysis on only those rotamers with sufficiently high-quality structural data supporting their conformations. Rather, we propose that, with the recent development of new structural refinement tools within Rosetta,[56, 57] these and other β-amino acid containing structures may be further refined, particularly now that there exist general, backbone-dependent rotamer libraries for the task.

### 2.4. Improved modeling of 3_14_ helix ligands

Encouraged by the agreement with higher levels of theory and the recovery of native rotamers, we undertook a retroactive binding affinity prediction. We rewrote a substantial part of Rosetta’s core libraries so that rotamer libraries might possess any number of dihedral angles and so that these rotamer libraries may be employed in the packer, Rosetta’s algorithm for concurrent sidechain optimization.

To test the ability of this new code and these new libraries to deliver scientifically useful results, we revisited an effort to design improved 3_14_ helix p53 mimetics from the Schepartz lab. Their data set included computational predictions via Monte Carlo free energy perturbation[58] (MC/FEP) in the OPLS-AA force field[59] as well as experimentally measured binding energies.[60] We sought to reproduce the trend in point mutation energies (ΔΔG) versus the base sequence. We compared the original MC/FEP result (*R*^2^ = 0.79) to a simple, standard ‘relaxation’ simulation carried out in Rosetta, using either the new or old rotamer libraries. Our rotamer libraries deliver comparable results (*R*^2^ = 0.80), without requiring any fit parameters. In contrast, with the prior rotamer libraries, which possessed the erroneous α-peptide like dependence on only cpand θ, performed quite poorly in comparison (*R*^2^ = 0.40) (Figure 6). This improvement suggests that a dependence on all three backbone angles is likely necessary for useful binding simulations.

**Figure 6:**
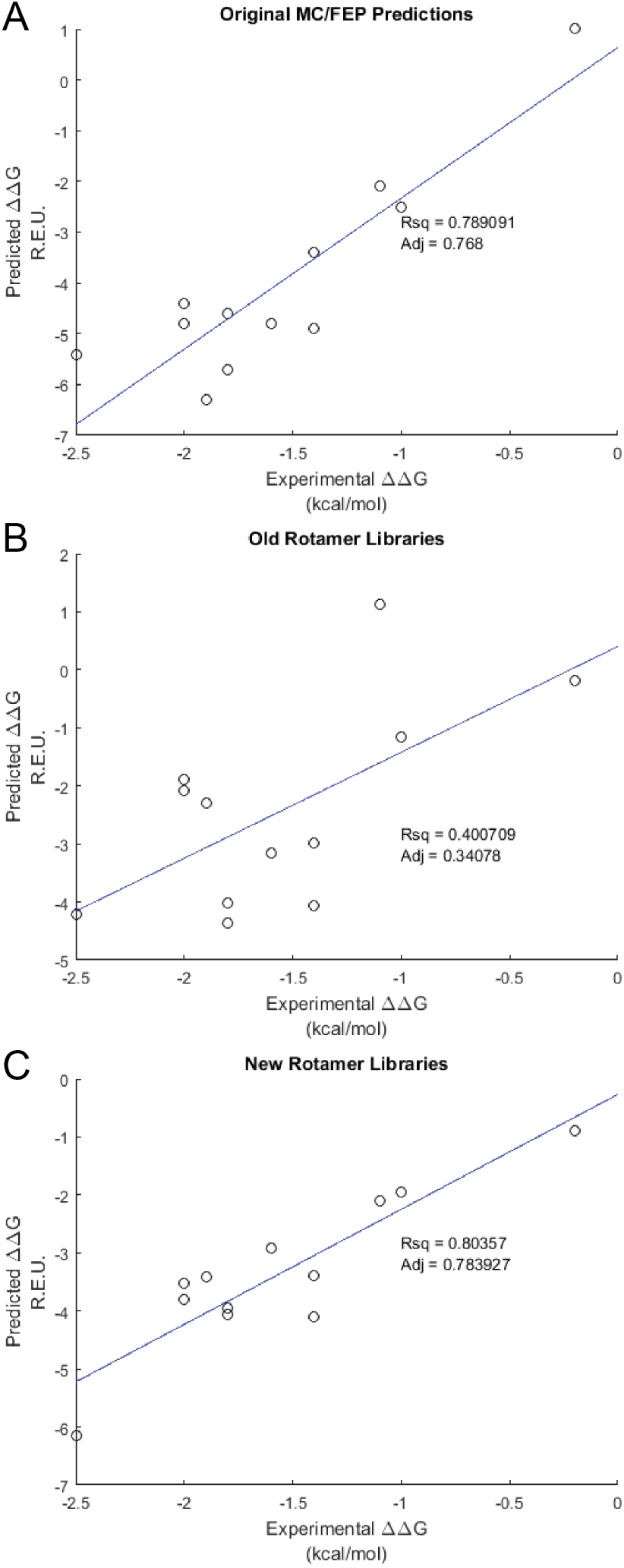
We compared experimental ΔΔG values (kcal/mol) for mutations in a series of 3_14_ helical p53 mimetics, which were assayed against the proteins mdm2 and mdm4, to computational predictions. The original predictions made by Schepartz (A) provided a standard of comparison; our Rosetta protocol performed poorly with the old (B) rotamer libraries but quite well using the rotamer libraries developed in this work (C).

## 3. Conclusion

We developed and tested a model of backbone-dependent libraries of β-amino acid side chain conformations. This model showed excellent agreement to simulations conducted at a higher level of theory as well as accurate recovery of experimental data, especially compared to previous models that borrowed α-amino acid conformations, without fitting additional scoring function parameters.

We incorporated new code and new residue parameter into the Rosetta software suite. Integration with Rosetta enables our contributions to open up new possibilities in design, modeling, and structure determination protocols. Furthermore, users of the software may parameterize new β-amino acid side chains of interest and obtain their rotamer libraries.

We anticipate eventual need for higher-resolution rotamer libraries (with more backbone bins, especially in highly populated regions). Doing so demands particular attention for β-amino acids because their backbone-dependent libraries are dramatically larger, both in memory and on disk. To this end, superior β-amino acid rotamer libraries will require more sophisticated interpolation methods.

Further efforts to create a framework suited for general heteropolymer modeling and design will include scoring function optimization to include alternative solvation and electrostatics models,[61, 62] reference energies for design, and better methods for modeling hydrogen and halogen bonds. Rerefinement of structures containing β-amino acids will provide an experimental basis against which that scoring function might be optimized. Taken together, these efforts will create the groundwork for designing arbitrary polymers of α- and β-amino acids.

## 4. Methods

### 4.1. β-peptide rotamer library creation

β-peptide rotamer libraries were created using the MakeRotLib protocol in Rosetta (Figure 7). For each backbone bin (from −170° to 180°, i.e.36^3^ = 46656 combinations), acetylated and methylamidated amino acid structures were initialized with varied side chain conformations. Residues with one χ (β^3^-Ser, β^3^-Thr, β^3^-Val) were sampled every degree; residues with two χ were sampled every 10° (1296 samples); residues with three (β^3^-Met; β^3^-Glu, β^3^-Gln) every 15° (13824 samples) or four (β^3^-Arg, β^3^-Lys) χ angles were sampled every 30° (20736 samples).

**Figure 7:**
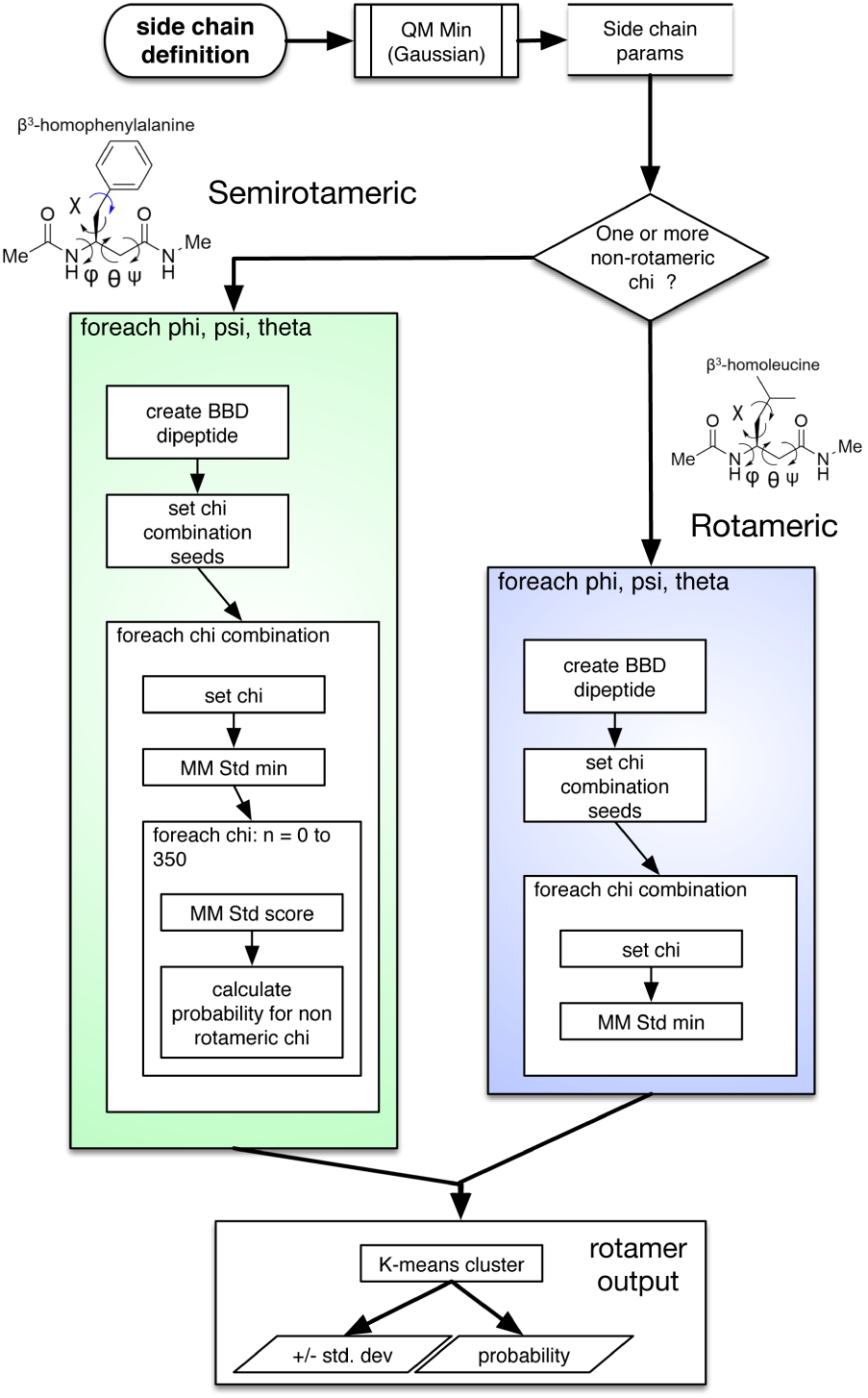
To construct rotamer libraries for β-amino acids, we create parameters for the residue by geometry optimization and charge fitting by QM as previously described. We seed backbone conformations, and for each one, we create hundreds or thousands of χ-combination seed conformations (omitting the final non-rotameric χfor semirotameric amino acids). We minimize each χcombination (and, for semirotameric residues, obtain a probability distribution for the terminal χ). Finally, we cluster, obtain a standard deviation in each dimension for each resulting rotamer well, and convert the minimized energies to probabilities via the Boltzmann distribution.

The resulting conformations were minimized using a linear, monotone minimizer and clustered using *k*-means clustering. The initial cluster centroids were chosen corresponding to physical intuition and to the choices made by the 2010 Dunbrack rotamer libraries.[12] Specifically, every rotameric χ was given a centroid initialized to g-, a, or g+, while nonrotameric χ angles had centroids every 30° (Symmetric nonrotameric χ angles such as aspartate’s second χ had six wells, 0° through 150°, while asymmetric nonrotameric χ angles such as asparagine’s second χ had twelve.) For each cluster, the energies for the rotamer sample closest to the cluster center are converted to Boltzmann factors and thereby to probabilities.

We also implemented functionality to provide a continuous treatment of terminal nonrotameric χ angles, as is possible in the 2010 Dunbrack libraries, thus permitting a smoother surface for energy minimization. In such rotamer libraries, created for β^3^D, β^3^E, β^3^F, β^3^H, β^3^N, β^3^Q, β^3^W, and β^3^Y, only the rotameric χ angles are clustered, while the final χ is sampled at 36 evenly spaced positions (5° increments for symmetric χ; 10° increments otherwise) and the energies in question are converted to probabilities similarly.

### 4.2. Evaluation of clash-free conformations

We sampled each amino acid at 5° intervals for every backbone dihedral angle. We minimized the sidechain degrees of freedom to ensure that the side chain was not making any unavoidable clashes. Then, we evaluated the distance between every pair of atoms and determined whether they were making a severe clash by comparing to 0.6 times the sum of the two atoms’ van der Waals radii. Thus, the value for α-amino acids represents a percentage, area-over-area, while for β-amino acids, the corresponding value is a percentage, volume-over-volume.

### 4.3. Structure optimization via Gaussian and comparison to the molecular mechanics scoring function

Using Gaussian09 revision D[63], we initialized acetylated and N-methylamidated structures of β^3^-Trp, β^3^-Leu, β^3^-Ile, and β^3^-Ser to two distinct and structurally valuable backbone conformations: (−140°, 60°, −120°), intended to be near stable conformations for the 3_14_ helix and the chimeric α3β-helix, and (65°, 75°, α140°), an extended state that can form pleated sheets like an α-amino acid forms β-sheets.

We initialized a 10° grid of side chain conformations (i.e. 1296 points per two-χ residue; 36 points for β^3^-Ser) and carried out energy optimization at the HF/6-311+G(2d,p) level of theory, followed by single point energy calculation at β^3^LYP/6-311+G(2d,p). Subsequently, we used OpenBabel[64] to generate PDB format files from the resulting Gaussian output and scored each structure using the molecular mechanics scoring function within Rosetta.

To analyze the similarity of the probability distributions, we evaluated the Bhattacharyya distance (Equation 1).

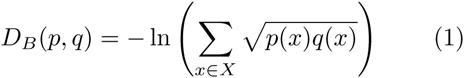

where *p* and *q* are two probability distributions and *X* is their domain.

The Bhattacharyya distance[65] is a generalization of the Mahalanobis distance that does not assume equal standard deviations; its range is [0, ∞), so values from 0.04 to 0.27 represent extremely similar distributions. Some deviations are, in fact, desirable and expected, as gas-phase quantum mechanics ought to deviate from molecular mechanics models that include solvation terms.

The worst fit is for the helical conformation of β^3^-Trp, where the MM scoring function appears to favor a β- χ_1_ dihedral. This result may partially result from Rosetta’s smaller van der Waals radii for hydrogen, as the g- rotamer may feature some minor clashes. The next worst fit is for β^3^-Ser, whose affinity for the g- rotamer in both backbone conformations relies in part upon a seven-membered intramolecular hydrogen bond that mm_std does not score and that the QM method may over-value due to the vacuum state. Importantly, even in the cases where there is disagreement, the best QM rotamers are always represented in the MM rotamer set and therefore will be sampled in simulations.

### 4.4. Comparison to PDB rotamers

For each crystal structure rotamer, we apply Rosetta’s tricubic interpolation to the nearest backbone bins to obtain an interpolated set of rotamers for the exact backbone dihedrals from the crystal structure. We measure the minimum RMS distance in degrees from a rotamer built for that position to the rotamer in the crystal structure and average that value for each residue type.

Figures from PDB structures are ray-traced in PyMOL. Electron density is generated from structure factors using phenix.maps or downloaded from the Uppsala Electron Density Server and visualized in PyMOL[66] using the isomesh command at two standard deviations within 1.6Åof atoms.

### 4.5. Rosetta simulations to predict binding affinity

We conducted simulations of the 3_14_ helices studied experimentally beginning with the same crystal structures of mdm2 and mdm4 as employed in the original computational study (PDB codes 2GV2 and 2Z5T, respectively). We generated point mutant structures of each 3_14_ helix ligand by mutating a position then repacking and minimizing an 8Å shell around the mutated residue.

We conducted a simple relaxation simulation, generating 1000 structures, to diversify the ensemble of β-peptide/protein complex structures, followed by interface scoring using the dGcross score term computed by the InterfaceAnalyzerMover. Separate sets of simulations were conducted with the old rotamer libraries (the canonical α-amino acid rotamers applied to the first two mainchain torsions) and with those developed in this work.

## 5. Supporting Information

### 5.1. Distinguishing features of β-amino acid rotamer libraries versus corresponding α-amino acid rotamer libraries

More than any other feature, rotamers for α- and β-amino acids are distinguished by steric clashes. As described throughout, many common rotamers compatible with an α-amino acid backbone are sterically incompatible with any β-amino acid backbone that shares φ and θ. A more minor influence is the absence of particular attractive interactions. For example, α-serine is well-known to form a hydrogen bond between its γ-H and its carbonyl oxygen in some backbone conformations, while that feature is largely absent in β^3^-serine: the six-membered hydrogen bonding ‘macrocycle’ is replaced with a much less favored seven-membered ‘ring’. In principle, β^2^-serine rotamer libraries would retain this backbone-dependent bias.

### 5.2. Comparison of β-amino acid rotamer recovery results to the performance of the Dunbrack libraries

It is challenging to directly compare two rotamer library models of different methodology and generality. This challenge, however, does not preclude the assessment of the fits of Figure 4. The Dunbrack libraries model rotamer wells as Gaussians with some standard deviation. We can employ these standard deviations to provide a natural scale for our observed average RMS errors. Let us assume that a Dunbrack rotamer is exactly centered on the “true” rotamer centroid for each χ angle. A typical standard deviation for such a Gaussian rotamer well is 8°. The expected RMS deviation from that rotamer well for one to four χ angles is 6.4°, 10°, 12.8°, and 15.1°. The standard deviation for this RMS deviation statistic is around 5° in each case. So, formally, the performance we observe reflects the same underlying variability of protein and peptide structures that the Dunbrack libraries do.

### 5.3. Detailed rotamer recovery for a structure of interest

To provide some context about the methods employed, we provide a worked example of rotamer recovery for two particular PDB structures: 4BPI, a complex between a Puma BH3 α/β peptide mimetic with Mcl-1, and 3G7A, an HIV gp41 bundle with an α/β peptide mimic of the CHR domain. Neither was an example of particularly good performance; rather, both illustrate the challenges of working with relatively sparse native structural data.

The former structure contains four β^3^-residues: β^3^D, β^3^R, β^3^Q, and β^3^E; while the latter contains two β^3^-homoglutamate residues. First, we had to revising atom and residue names in Rosetta to match Rosetta’s nomenclature, to make sure that no atoms were wrongly detected as missing, which would have resulted in coordinates being built from ideal. In fact, we discard the β^3^E residue from 4BPI on these grounds, as it is involved in binding a cadmium atom in the native structure, and one of the β^3^E residues from 3G7A, as its Cγ and Oδ atoms are missing. Second, we built rotamer sets for each residue. The closest interpolated rotamers to each experimental rotamer are given in Table S1.

**Table S1:** Example recovery of native rotamers; unusual native rotamers are not well recovered.

| PDB | Residue | Native | Interpolated |
| --- | --- | --- | --- |
| 4BPI | $\beta^3$D | (-2.342, -19.669) | (-21.486, -21.272) |
| 4BPI | $\beta^3$R | (-77.714, -149.718, -0.905, 102.730) | (-80.805, -112.957, 1.527, 66.799) |
| 4BPI | $\beta^3$Q | (-89.278, -102.220, 165.450) | (-94.279, -86.110, 159.207) |
| 3G7A | $\beta^3$E | (-65.248, 142.906, 78.350) | (-57.471, 164.515, 82.988) |

Native conformations so far from the theoretical sp^3^-sp^3^ optimum, especially in mostly surface-exposed positions, seem surprising. Reproducing them perfectly would be an inappropriate burden for a rotamer library; this library’s attempts to do so largely hinge on very low-probability rotamers that would likely be rebuilt in most simulations. Indeed, their distance from optima suggests that they could reflect structural errors that remain uncorrected simply because of the limited knowledge surrounding the preferred side chain conformations of β-amino acids.

The B-factors for each β^3^ residue in these structures are quite high, from 45 to 70. Examining maps derived from the deposited structure factors, experimental evidence for these particular conformations seems either weak or absent entirely (Figure 5). Ironically, more side-chain electron density is present for the lower-resolution structure (3G7A, at 2.8Å, at least able to secure the relative position of the α and γ carbon atoms) than for the higher-resolution structure (4BPI, at 1.98Å).

### 5.4. Code availability

Applications to create amino acid rotamer parameter files and rotamer libraries, as well as the resulting rotamer libraries are distributed as part of the Rosetta software suite, which is free for academic use.

## 6. Acknowledgements

A.M.W. is supported by a NYU Dean’s Dissertation Fellowship. P.S.A. thanks the National Institutes of Health (R01GM073943) for financial support of this work. R.B. and P.D.R are supported by the Simons Foundation, Center for Computational Biology.

